# Discovery, induction, and screening of prophages in clinical *Acinetobacter baumannii* isolates

**DOI:** 10.64898/2026.02.16.706188

**Authors:** Jessica Trinh, Vivek K. Mutalik, Catherine M. Mageeney

## Abstract

**Background:** *Acinetobacter baumannii* is a common bacterial pathogen in nosocomial infections. It has become one of the greatest threats to human health for its growing resistance to ‘last resort’ antibiotics, which has led to a revival of phage therapy as a potential treatment. However, conventional methods for isolating *A. baumannii*-infecting phages are labor-intensive and often unsuccessful.

**Methods:** Our approach involves a computational pipeline to identify temperate phages (prophages) integrated into *A. baumannii* genomes, followed by mitomycin C (MMC) induction of those strains to screen for active prophages.

**Results:** Here we show a prophage analysis for nearly 900 *A. baumannii* genomes. We observed MMC-triggered excision of nine prophages from eight *A. baumannii* strains by PCR and sequencing. Further we show four prophage form virions detectable by transmission electron microscopy, and two which can plaque on other *A. baumannii* isolates.

**Conclusion:** This work demonstrates the utility and diversity of prophages for further development as therapeutics for antibiotic resistant *A. baumannii*.

## Introduction

Antibiotic resistance (AR) is a significant threat to global public health; 4.95 million deaths across the world were estimated to be associated with bacterial AR in 2019 (Antimicrobial Resistance Collaborators 2022), highlighting the need for new therapeutics against AR pathogens. The six ‘ESKAPE’ organisms (*Enterococcus faecium, Staphylococcus aureus, Klebsiella pneumoniae, Acinetobacter baumannii, Pseudomonas aeruginosa*, and *Enterobacter sp*.) are of particular interest due to their alarming rise in antibiotic resistance and high rates of infection. These organisms display natural resilience to survive for extended periods of time under wide range of environmental conditions and are the cause of frequent infection outbreaks in hospitals (Miller and Arias 2024). Carbapenem-resistant *A. baumannii* in particular is ranked as a critical-priority pathogen by the World Health Organization based on factors such as limited treatability, prevalence, and infection mortality (Willyard 2017, Sati *et al*. 2025). In some countries, rates of carbapenem-resistance in *A. baumannii* can be as high as 80 – 90% (Miller and Arias 2024).

Bacteriophages (phages) are the natural predators of bacteria that have been studied extensively for longer than a century (Altamirano and Barr 2019). AR has led to a revival of leveraging phages as therapeutic agents against resistant infections (“phage therapy”, Chan *et al*. 2018, Jennes *et al*. 2017). Across the world multiple cases have been treated using phage therapy for different antibiotic-resistant bacterial pathogens, including *A. baumannii* (Altamirano and Barr 2019, Liu *et al*. 2022, Schooley *et al*. 2017, Wang *et al*. 2025). Despite these successes, phage therapy faces significant implementation challenges. Phages often exhibit narrow host ranges with strain-level specificity, making it challenging to find phages for specific target strains. Additionally, phages are typically isolated by screening environmental samples such as raw sewage or soil, against patient isolates. Though there are currently ongoing efforts to build phage banks for high priority bacterial pathogens, this process becomes resource and time-intensive for some clinical isolates. In some instances, no phages are recovered even after multiple isolation attempts from patient, environmental, or animal samples Goh *et al*. 2005, Meesen-Pinard *et al*. 2012).

To address the limitations of traditional phage discovery, researchers have investigated the use of prophages as potential therapeutic agents. Prophages are phages integrated into bacterial genomes and are found in nearly every bacterial species (Dieppa-Colon *et al*. 2025, Yu *et al*. 2025, López-Leal *et al*. 2022). They are typically overlooked as therapeutics due to concerns about transduction, lysogenic conversion, and the potential to increase virulence through horizontal gene transfer (Cook *et al*. 2025, Monteiro *et al*. 2019, Strathdee *et al*. 2023). However, prophages can be converted into virulent phage through genome engineering that removes integration machinery and other potentially deleterious gene products (Mageeney *et al*. 2020b, Kakkar *et al*. 2024, Dedrick *et al*. 2019). This approach provides researchers with an alternate method to access a large array of phages when traditional phage discovery is unsuccessful.

In this study, we employed the Phage Factory platform (Mageeney *et al*. 2020b) to identify prophages within the genomes of hundreds of *A. baumannii* strains. We identified 789 prophages across 891 *A. baumannii* strains. Comparison of this dataset to previously isolated *A. baumannii* phages deposited in GenBank, revealed increased diversity suggesting the prophages provide an expanded pool of phages for therapeutic purposes. We then chose eight strains to understand prophage kinetics and host ranges. Following mitomycin C (MMC) induction, we verified nine actively excising prophages through PCR and deep sequencing analysis. Analysis of the induced lysates revealed that four strains contained prophages capable of producing virion particles, and two of these produced phages capable of plaquing on strains in the collection. Our study expands the repertoire of *A. baumannii*-infecting phages with potential therapeutic applications and provides insights into prophage replication dynamics.

## Results

### Prophages are abundant in A. baumannii isolates

Our prophage discovery pipeline uses two programs to identify integrative genomic elements (IGEs), which includes prophages. TIGER uses a comparative genomics approach to identify IGEs that contain either tyrosine or serine integrases (Mageeney *et al*. 2020a, Mageeney *et al*. 2022, Yu *et al*. 2025), while Islander detects IGEs encoding a tyrosine integrase which target tRNA or tmRNA genes for integration (Hudson *et al*. 2015). We analyzed a curated set of 891 *A. baumannii* genomes, including 100 strains from a recently isolated panel of diverse strains (Galac *et al*. 2020, **Supplemental Table S1**). TIGER and Islander predicted 789 total prophages among all tested strains, with individual strains containing zero to five prophages (**Figure 1, Supplemental Table S2**). Approximately one-third of strains contained no prophages (323/891, 36.3%), while only 18.1% harbored two or more prophages (161/891). TIGER and Islander also identified integration sites (bacterial attachment site; *attB*) for putative prophages. Analysis of unique *attB* sites revealed that more than half of the predicted prophages integrated into tRNAs and other non-coding RNAs (51.3%), while the remaining integrate into protein coding genes (48.7%, **Figure 2**).

**Figure 1.**
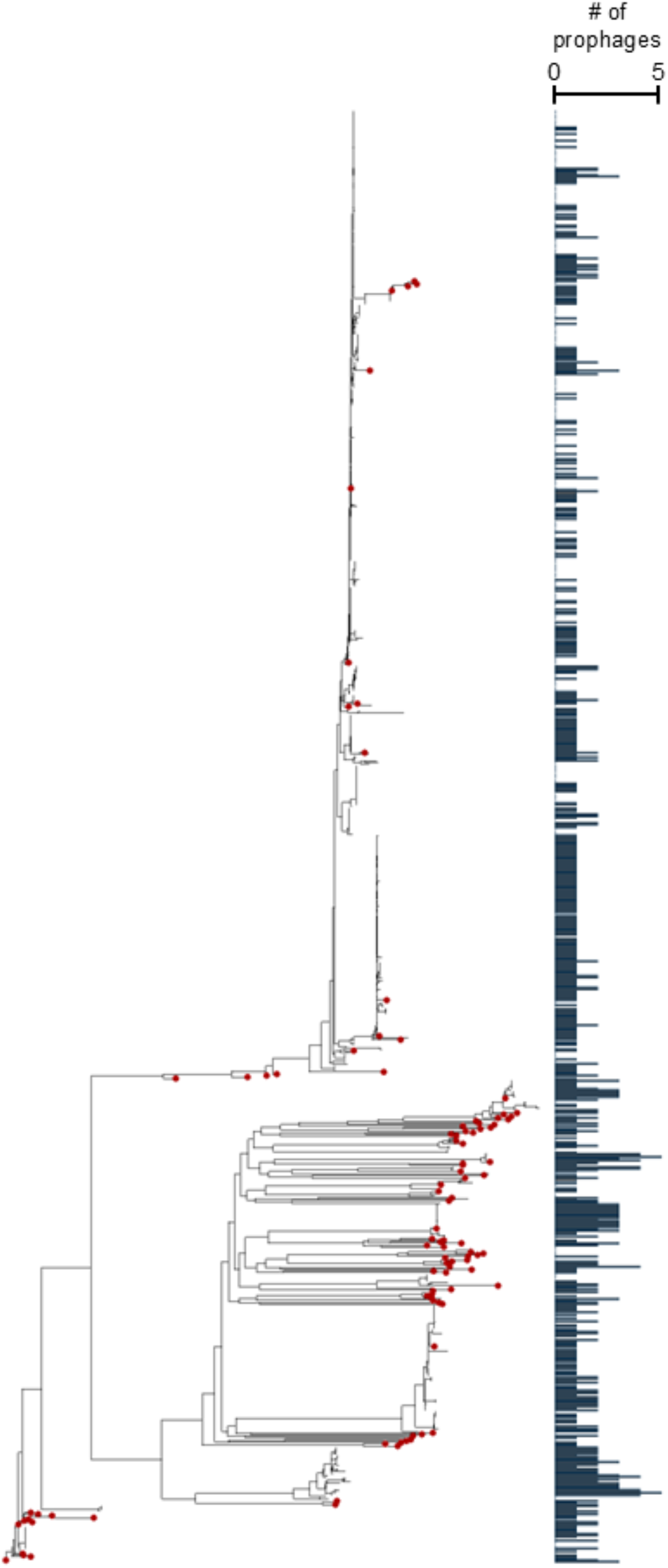
*A. baumannii* strains form distinct phylogenetic groups, with varying amounts of putative prophages in each genome. Strains are plotted on a mid-rooted core genome tree to represent the diversity of the 861-genome data set. Bar plot (right) indicates the number of prophages each strain has from zero to five prophages. Red dots indicate strains from the Galac *et al*. 2020 dataset.

**Figure 2.**
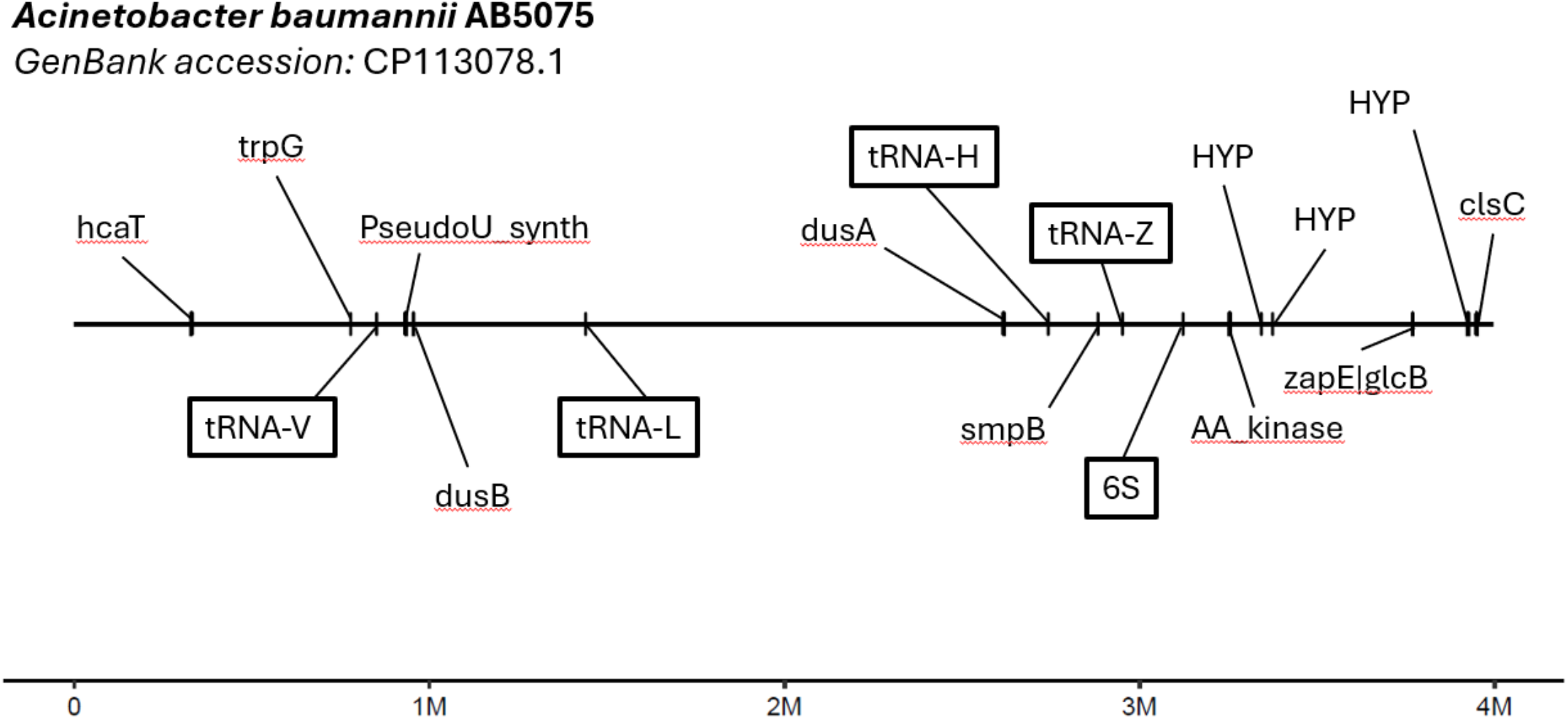
Chromosome map featuring unique integration sites (attB) for predicted prophages, superimposed onto the genome assembly of *A. baumannii* AB5075 (GenBank accession: CP113078.1). Integration sites are labeled with the region that the site corresponds to, with boxed labels indicating sites of non-protein-coding features such as tRNAs and rRNAs.

To assess the diversity of the predicted prophages and determine whether they expand the current phage repertoire for *A. baumannii*, we clustered them with 508 *A. baumannii* deposited in NCBI GenBank. Clustering analysis revealed 42 total clusters, of which 14 contained prophages. The TIGER/Islander-predicted prophages tend to cluster together, with most exhibiting mosaic genomic architecture (**Figure 3**). Some phages that clustered with prophages lacked an annotated integrase. However, additional analysis identified recombinases and/or repressor proteins in these sequences (see Methods; **Supplemental Table S3**). These predicted prophages substantially increase the known diversity of *A. baumannii*-infecting phages beyond what has been discovered through traditional isolation methods.

**Figure 3.**
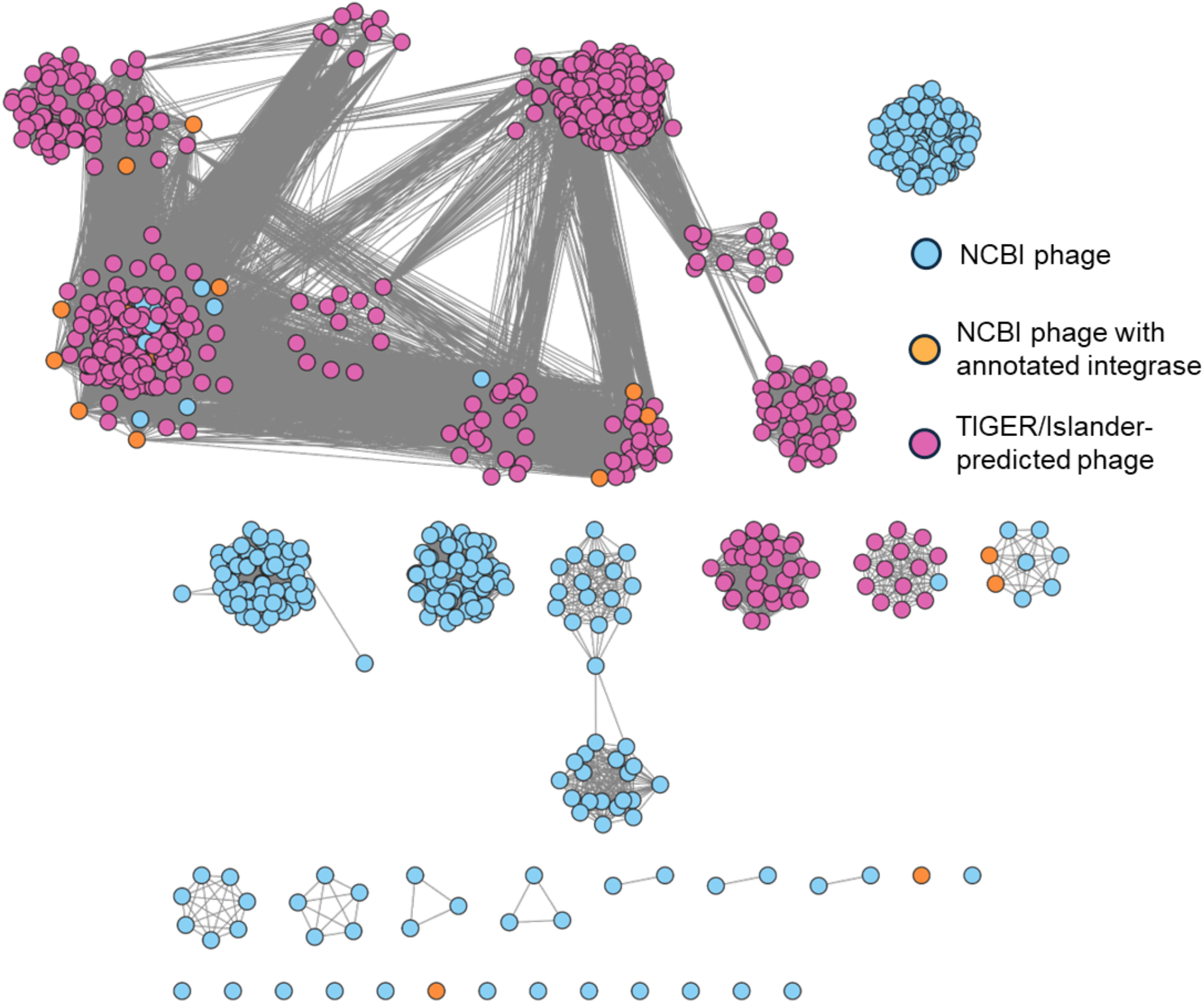
Prophages in *A. baumannii* form many related clusters. Clusters are computed via Markov Cluster Algorithm using percent amino acid identity and percent shared genes (Nayfach *et al*. 2021). TIGER/Islander-predicted prophages have purple labels. NCBI phages that contain any gene already labeled as an “integrase” have orange labels. All other NCBI phages are labeled blue. Clusters were used to determine prophage candidates for further screening by eliminating candidates that were likely to be identical or highly similar.

### MMC induction reveals active prophages from nine different strains with variable induction kinetics

We selected eleven strains for further analysis of prophage induction and activity based on the following criteria. The predicted prophages needed to be 40 to 60 kbp in size, contain a complete set of genes expected for functional phages (including terminase, capsid, tail, portal, and lysin proteins), and represent different clusters.

A hallmark of MMC-induced prophage replication is a decrease in OD several hours after MMC treatment (Humphrey *et al*. 1995, Filipiak *et al*. 2020). Eight of the eleven strains tested showed a decrease in OD after at least 4 hours of MMC treatment (**Supplemental Figure 1**). Because TIGER and Islander precisely predict prophage boundaries, we were able to amplify the phage attachment (*attP*) site, a recombination site formed by circularization of the excised prophage, for nine prophages across the eight strains (**Figure 4**). Only one predicted prophage failed to show evidence of excision through *attP* site detection (2575.42.V.x).

**Figure 4.**
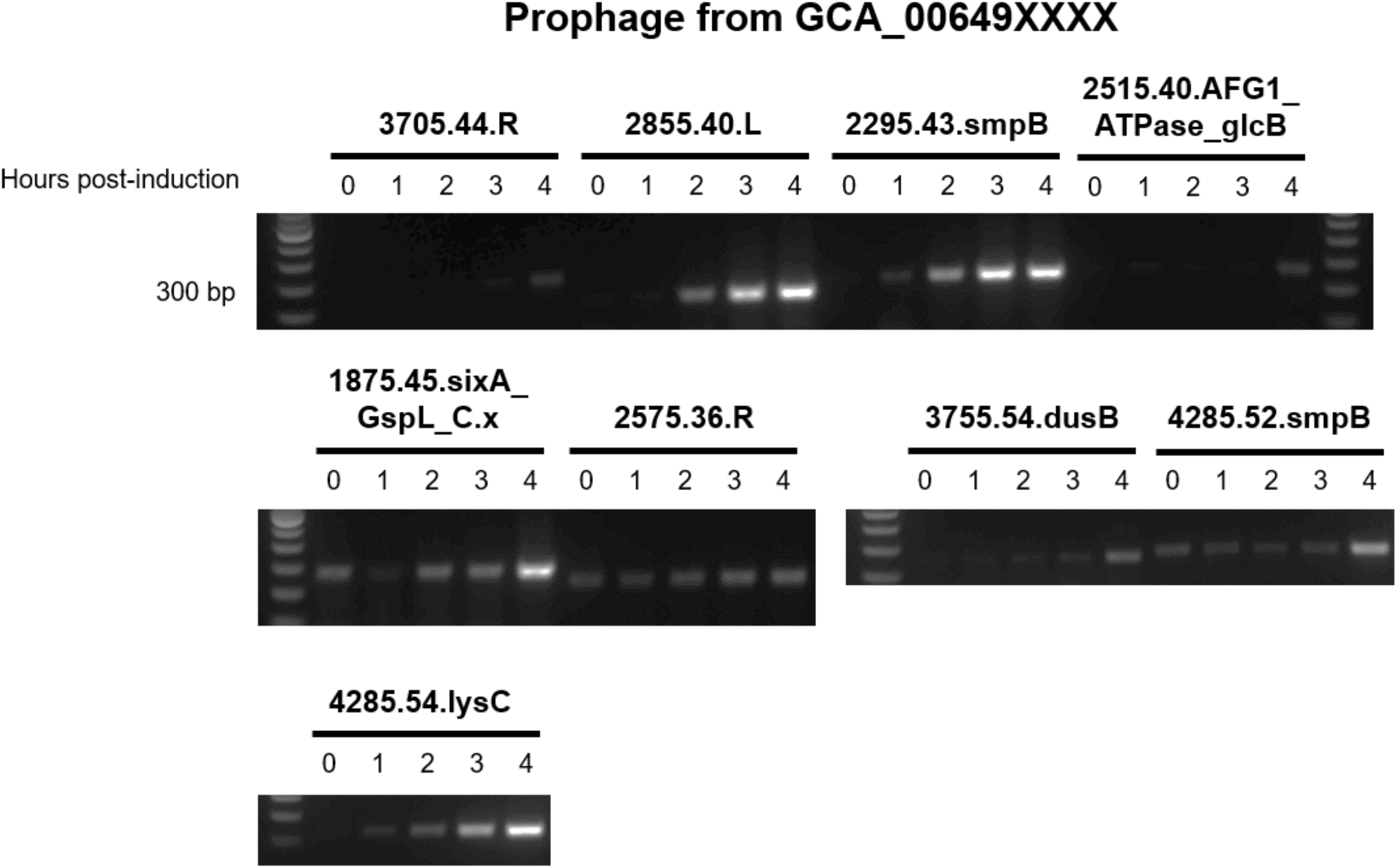
Detection of TIGER-predicted prophage in *A. baumannii* supernatants. 1% agarose gels showing PCR products (200-300 base pairs) amplifying the *attP* site from DNase-treated *A. baumannii* lysates. Prophages are named after the last 4 digits of their GCA accession number, the size of the prophage in kilobases, and their integration site. A positive *attP* site band suggests the presence of actively-replicating prophage. Note that one strain has more than one prophage detected by PCR (GCA_006494285).

We spot tested lysates from these strains on nineteen *A. baumannii* strains including the eleven prophage-laden strains from our initial set plus eight prophage-free close relatives). MMC-induced lysates from two strains produced plaques (**Supplemental Figure 2**). Interestingly, both strains infected by these induced prophages themselves contain at least one putative prophage. Typically, prophage containing strains prevent secondary phage infections through superinfection exclusion, usually through a prophage-encoded repressor proteins (Heo *et al*. 2007, Bondy-Denomy *et al*. 2016, Dedrick *et al*. 2017).

We further characterized the induction biology of these prophages by deep sequencing the MMC-induced genomic DNA to reveal excision and replication kinetics for the nine phages active identified in our PCR screen (**Figure 5**). This method allowed us to determine differences in induction timing and replication rates for each prophage, as well as validate additional putative prophages in the same genomes that were originally excluded by during selection.

**Figure 5.**
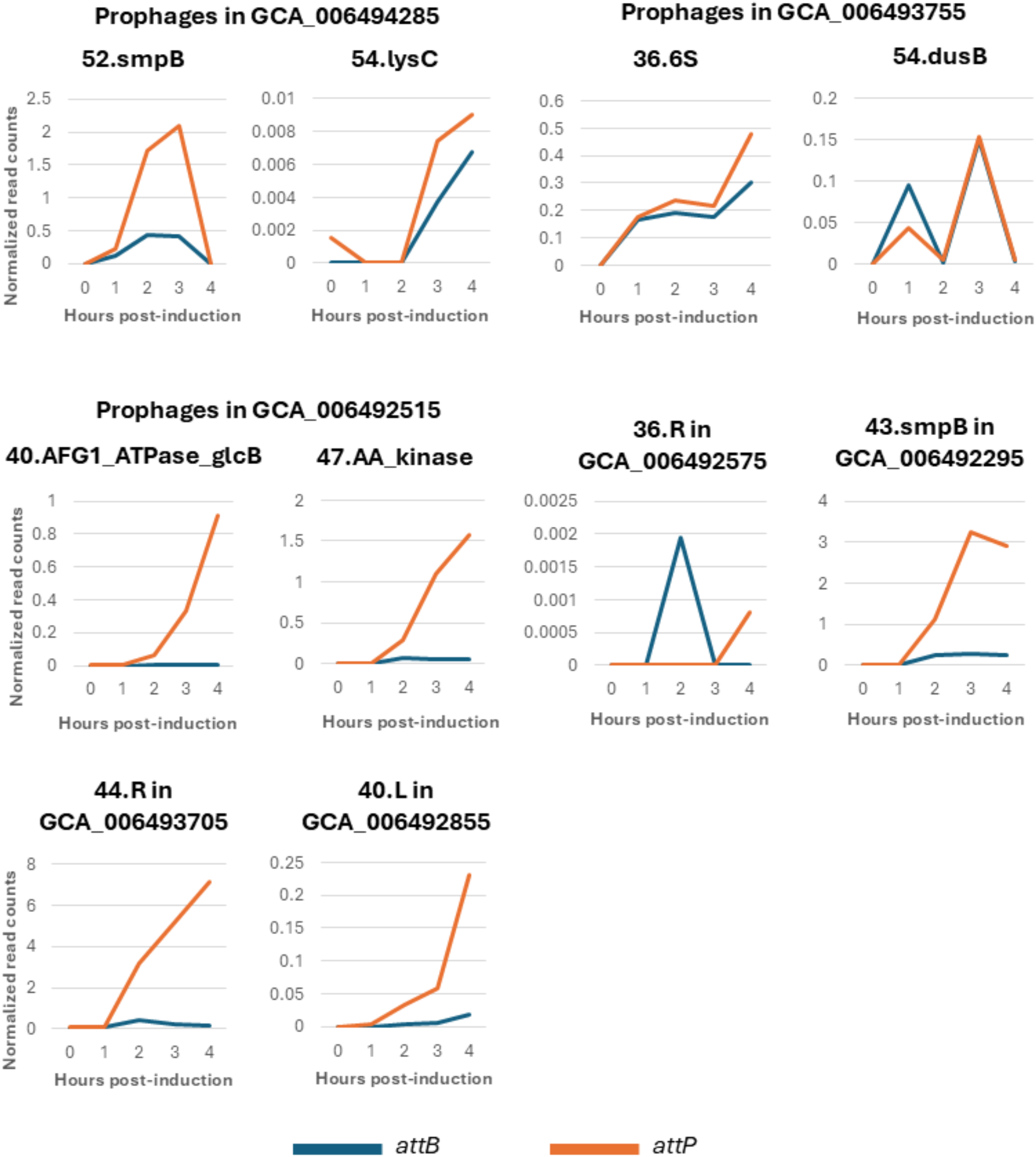
Short-read sequencing data reveals that prophages in *A. baumannii* undergo different induction dynamics. Line graphs depict the normalized abundances of reads that map to *attB* (junction left after prophage excision) versus those that map to *attP* (excised circularized phage genomes). Read counts are normalized to the quantity of reads that map to *attL* and *attR*. Note that y-axes scales are different for each graph, and some genome assemblies contain more than one prophage.

Sequencing data confirmed that both predicted prophages in strain GCA_006494285 (52.smpB and 54.lysC) underwent excision but their induction timing is differed. Prophage 52.smpB induced early with peak excision at three hours post induction, while 54.lysC induced more slowly and began peaking at four hours post-induction. Additionally, reads mapping to the *attP* site of 52.smpB were four times more abundant that reads mapping to its corresponding *attB* site at three hours post-induction, which indicates phage replication. Similar trends were observed for strains that only contain one prophage, with phage genome replication verified to occur between two- and four-hours post-induction (**Figure 5**). While the PCR results confirmed that the predicted prophages were actively excising, sequencing data revealed that the intensity and timing of induction varied among different phages.

### Transmission electron microscopy reveals phage particles in induced A. baumannii supernatants

Many of the predicted prophages showed genomic evidence of excision but only two produced plaques. To verify whether these prophages could produce functional virions we used transmission electron microscopy (TEM) to visualize phage particles in the MMC-induced lysates. We tested eight MMC-induced lysates and detected intact phage particles in four of them (**Figure 6**). All particle-containing lysates contained phages with icosahedral heads. Phages in the supernatants of strains GCA_006493705, GCA_006491875, and GCA_006492855 had consistent capsid diameters ranging from 53 to 70 nm and tail fiber lengths ranging from 162 to 302 nm (**Supplemental Table S7**). Notably, lysates from two strains (GCA_006492575 and GCA_006492855) contained phage particles with unique morphologies. Virions from GCA_006492575 possessed tails with unusual protein structures attached (**Figure 6C**), while GCA_006492855 contained multiple phages with round capsids attached to long, thin pilus-like structures (**Figure 6D**). Unfortunately, these phages could not be propagated from single plaques, making them difficult to isolate.

**Figure 6.**
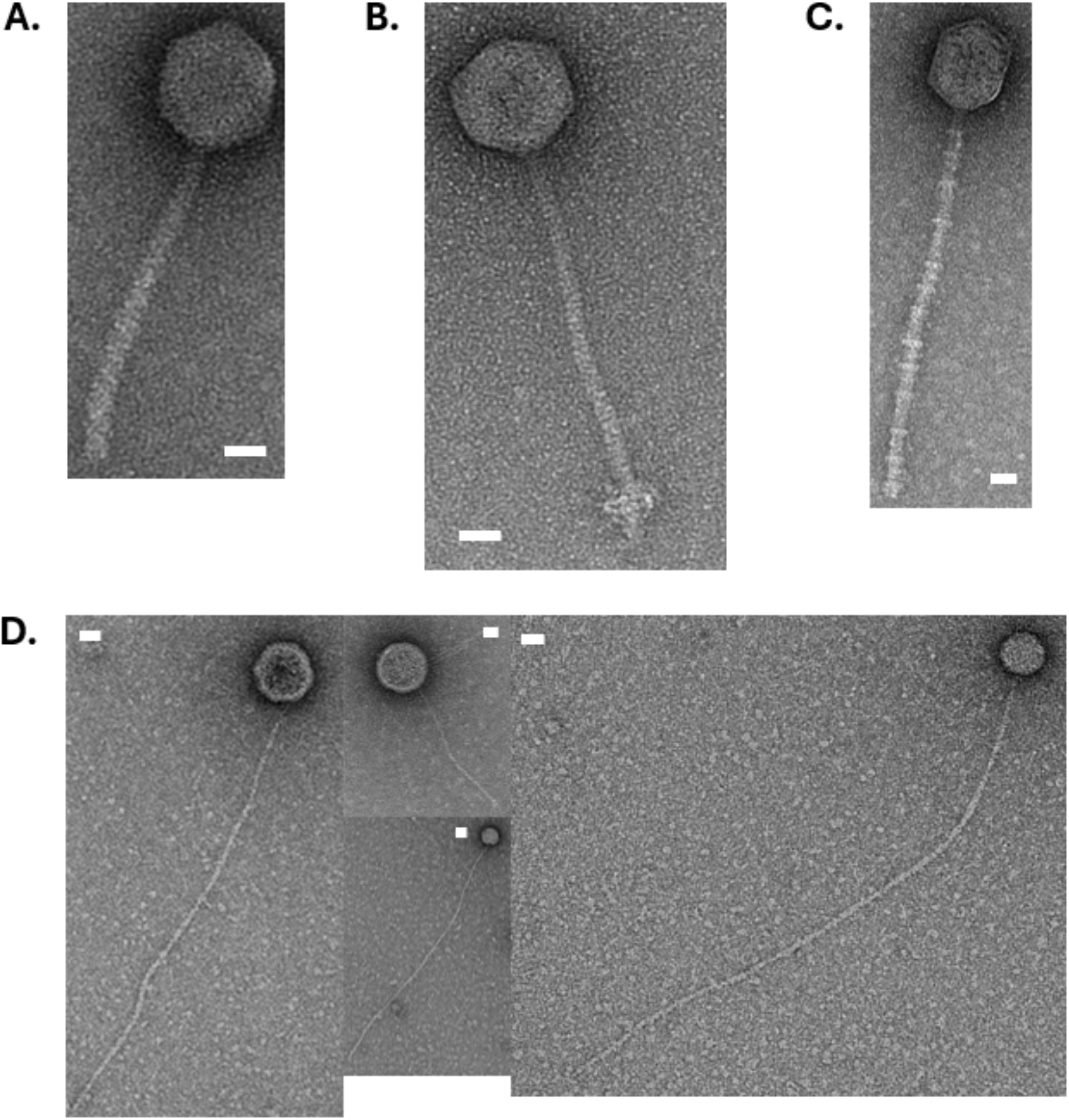
Visualization of phage particles from supernatants from *A. baumannii* strains using transmission electron microscopy. Phage particle(s) in each panel are from supernatants of the following strains: A. GCA_006493705, B. GCA_006491875, C. GCA_006492855, D. GCA_006492575. White scale bar = 20 nm. Note that lysate from D contains multiple morphologies of phage particles.

## Discussion

In this study, we used computational tools to predict prophage within *A. baumannii* genomes and characterized their activity using culture-based and molecular-based methods. While other studies have examined the diversity of prophages within *A. baumannii* (Costa *et al*. 2018, Tenorio-Carnalla *et al*. 2024, Loh *et al*. 2020, López-Leal *et al*. 2020), relatively few temperate phages have been evaluated for therapeutic use (Peng *et al*. 2022, Mardiana *et al*. 2023). Furthermore, very few studies investigated the dynamics of MMC-induced prophage activation (Mageeney *et al*. 2020b, Donovan *et al*. 2015). Here, we precisely predicted multiple prophages from *A. baumannii* that are unique compared to isolated phages deposited in GenBank. We have further characterized phage excision, replication, and virion production using multiple assays to determine if prophages would be useful as potential therapeutic agents. Our analysis of prophage population dynamics in strains containing one or two prophages reveals the complexity of prophage excision timing and replication kinetics.

While other recent efforts have predicted prophages (Costa *et al*. 2018, Tenorio-Carnalla *et al*. 2024, Loh *et al*. 2020, López-Leal *et al*. 2020), these studies typically used gene content analysis methods that cannot precisely characterize the attachment sites for individual prophages. Our methods provide this precision, enabling detailed investigation into infection kinetics, polylysogeny dynamics, and prophage activity. We demonstrate that 84.6% (11/13) of predicted prophages from our selected strains were capable of genomic excision. Of these, four produced enough phage particles for TEM visualization, and two produced plaques on strains within our collection. Data from our deep sequencing analysis highlights the diversity of induction kinetics across strains and even within the same strain. These findings are essential for evaluating the therapeutic potential of prophages, either from their native temperate form where bacterial lysis occurs upon stress induction or as engineered virulent derivatives.

To use prophages as therapeutic agents, they must be engineered into virulent phages by removing integrases, repressors, and operator sequences (Mageeney *et al*. 2020b, Kakkar *et al*. 2024, Dedrick *et al*. 2017, Kilcher *et al*. 2018), and then further characterized *in vivo* for safety and efficacy (Pires *et al*. 2020, Mutalik and Arkin 2022). Compared to other Gram-negative bacteria, *A. baumannii* presents two major challenges for effective phage engineering. First, even though CRISPR-based phage engineering technologies have been established in many other Gram-negative bacteria (Kiro *et al*. 2014, Hupfield *et al*. 2018, Chen *et al*. 2019), such methods in *A. baumannii*, are still being developed (Wang *et al*. 2022, Wang *et al*. 2019). Additionally, highly efficient CRISPR enzymes remain limited in this species. Secondly, while it is possible for phage to be engineered through synthesis and/or transformation of engineered phage genomic DNA (also known as phage “rebooting”, Ando *et al*. 2015, Kristensen *et al*. 2024, Ipoutcha *et al*. 2024), very few *A. baumannii* phages have been successfully rebooted in model systems like *E. coli* and have shown reduced efficiency compared to other Gram-negative phages attempted (Cheng *et al*. 2022). This highlights the need to understand which phage features enable successfully rebooting. As phage therapy advances, developing methods for engineering and rebooting in non-model systems becomes increasingly important. While further research is needed to successfully engineer and deploy prophages as therapeutics, we demonstrate the increased diversity these prophages provide and their potential utility for phage therapy applications.

## Materials and Methods

### Prophage prediction, clustering, and host strain phylogeny in Acinetobacter baumannii

Prophage prediction was performed as described previously (Mageeney *et al*., 2020b). Briefly, two discovery platforms for IGE, Islander (Hudson *et al*., 2015) and TIGER (Mageeney *et al*., 2020a) were applied to 891 *A. baumannii* genomes from GenBank. Genomic islands were annotated with our Tater software (Mageeney *et al*. 2020a). The software uses Prodigal to call open reading frames, Prokka and Pfam-A HMM database were used to prediction phage protein functions. IGEs are sorted into different categories based on the type of island. 789 phages were identified as “Phage 1”, which are prophages with a credibly complete genome, while 289 phages were identified as “Phage 2” because they may be missing some phage genes. The software detected only five putative filamentous phage. All candidates identified as “Phage 1” are further filtered based on genome size and whether these phages are present in unique clusters.

A core genome tree for all *A. baumannii* strains was built with IQ-TREE (v.3.0.1, Minh *et al*. 2020) using the ModelFinder algorithm (Kalyaanamoorthy *et al*. 2017) to determine the appropriate substitution model and 1,000 ultrafast bootstrap replicates to generate a consensus tree. Prophage counts were mapped onto the resulting phylogenetic tree using the R package ggtree (v.3.10.0, Yu *et al*. 2017). Prophage DNA sequences were then clustered by Markov Cluster Algorithm based on an average amino acid identity (>40%) and minimum percent of genes shared (>20%, Nayfach *et al*. 2021, https://github.com/snayfach/MGV/blob/master/aai_cluster/README.md) to select phages for screening by minimizing the number of phages in the same cluster screened. Phages from NCBI were quickly screened for integrases with a keyword search, and other phages from NCBI that clustered with TIGER/Islander-predicted prophages were evaluated further with HMMER and HMMs from the PHROGs dataset (https://www.ebi.ac.uk/Tools/hmmer/search/phmmer, Terzian *et al*. 2021) for integrases and repressor proteins.

### Prophage induction and plaque assays

For the initial screening of prophage-laden strains, *A. baumannii* strains were grown overnight at 37°C in LB broth and back-diluted 1:100 into 150 uL LB in sterile 96-well plate wells (Fisher Scientific #08-772-54). Optical density was measured every 10 minutes on a Tecan Spark plate reader. When cultures reached an OD600 of 0.4 to 0.5, cells were induced with 1 μg/mL MMC and incubated for up to six hours (**Supplemental Figure S1**). Three technical replicates were used for each strain, and graphs were generated in Microsoft Excel showing the average of each well per timepoint measured. Eight prophage-containing strains with selected prophages were selected for further analysis.

To collect lysates and genomic DNA, *A. baumannii* strains were grown overnight at 37°C in LB broth, back-diluted 1:100 in 5 mL LB, and incubated to an OD_600_ of 0.4 to 0.5. Cultures were then induced with 1 μg/mL MMC (Fisher #BP25312). 1 mL aliquots were collected from cultures at 0, 1, 2, 3, and 4 hours post-induction. Cells from aliquots were pelleted by centrifugation at 6,000 × *g* for 10 minutes. Supernatants were harvested and filtered through a 0.2-μm syringe filters (Whatman^®^ #WHA9916-1302) while cell pellets were stored at -20°C for subsequent genomic DNA extractions. Plaques were generated either by mixing filtered supernatants with 4 mL of 0.6% (soft) agar and 20 μL of host bacterial culture or by spotting 3 μL supernatant onto host lawns made with 4 mL soft agar and 20 μL of host bacterial culture. Plates were incubated overnight at 37°C to visualize plaques on host lawns.

### Assessing prophage activity with PCR and deep sequencing

One output of the TIGER program is a set of predicted *att* sites indicating the recombination sites of the integrated or replicating prophage (*attL, attR, attB, attP*, Mageeney *et al*. 2020a). Primers to test for actively replicating phage were designed according to the *att* sites computed for each prophage (**Supplemental Table S5**). Supernatants from prophage induction treatments were then DNaseI-treated (Norgen Biotek #25710) to remove DNA not protected by capsids and 2 μL was used as template for PCR detection of *attB* and *attP* sites, indicative of excised and circularized prophage genomes. To explore prophage dynamics we used deep sequencing. Genomic DNA was isolated from cell pellets using the Qiagen DNeasy blood and tissue kit (Qiagen #69504). Sequencing libraries from genomic DNA were prepared using the Illumina DNA Prep, (M) Tagmentation kit (Illumina #20060059) and using Illumina® DNA/RNA UD Indexes Set B (Illumina #20091656) following the manufacturer’s recommended protocol.

DNA libraries were quantified using the Qubit High-Sensitivity dsDNA Quantitation kit (ThermoFisher #Q32851) and pooled in equal quantity to make a final library. Quality control on the final library was performed with the Qubit quantitation kit and 4150 TapeStation System (Agilent #G2992AA) using the HighSensitivity D5000 Reagents and ScreenTape (Agilent #5067-5592 and #5067-5593). Samples were either shipped to Genewiz for Illumina sequencing or sequenced in-house on a NextSeq™ 2000 (Illumina) system using the NextSeq™ 2000 P3 XLEAP-SBS™ Reagent Kit (200 Cycles) (cat # 20100989), generating an average of 15 million reads per sample to ensure quantification of low-abundance reads. Reads were processed through Juxtaposer (Schoeniger *et al*. 2016), a software package built to detect genomic recombination in sequencing reads. We generated probes for *attB, attP, attL*, and *attR* to detect and count recombinant reads (**Supplemental Table S6)**.

### Transmission electron microscopy imaging of phage

Supernatants from induced bacterial cultures were concentrated by spinning samples down at 5,000 RPM for 4 hours at 4°C before shipping 50 μL of the supernatant to the University of Maryland, Baltimore County (UMCB) for further analysis. Imaging work was performed by Tagide deCarvalho at the Keith R. Porter Imaging Facility at UMBC on a Hitachi HT7800. Capsid diameters and tail lengths were measured with ImageJ software using a provided scale bar as a reference.

## Authors’ Contributions

Conception and design of the work were contributed by J.T., V.K.M., and C.M.M.. J.T. executed experiments and performed most of the bioinformatic analyses. C.M.M supervised the work and supported bioinformatic analyses. *A. baumannii s*trains and genome assemblies were curated and provided by V.K.M. J.T. and C.M.M. wrote the manuscript. All authors read, commented on, and approved the final article.

## Acknowledgements

We thank Alexey E. Kazakov (Lawrence Berkeley National Laboratory) for assisting with the curation of *A. baumannii* genomes for this project. We also thank Kelly P. Williams and Ellis L. Torrance, and Hannah McClain (Sandia National Laboratories) for their work on the Phage Factory pipeline, Juxtaposer, and *att* site software (manuscript in prep). We thank Joanna Steczynska (Sandia National Laboratories) for her thoughtful feedback on this manuscript.

Sandia National Laboratories is a multi-mission laboratory managed and operated by National Technology & Engineering Solutions of Sandia, LLC (NTESS), a wholly owned subsidiary of Honeywell International Inc., for the U.S. Department of Energy’s National Nuclear Security Administration (DOE/NNSA) under contract DE-NA0003525. This written work is authored by an employee of NTESS. The employee, not NTESS, owns the right, title and interest in and to the written work and is responsible for its contents. Any subjective views or opinions that might be expressed in the written work do not necessarily represent the views of the U.S. Government. The publisher acknowledges that the U.S. Government retains a non-exclusive, paid-up, irrevocable, world-wide license to publish or reproduce the published form of this written work or allow others to do so, for U.S. Government purposes. The DOE will provide public access to results of federally sponsored research in accordance with the DOE Public Access Plan.

## Conflict of Interest

The authors declare no potential conflicts of interest.

## Funding Statement

This was completed as part of the BRaVE Phage Foundry which is supported by the U.S. Department of Energy, Office of Science, Office of Biological & Environmental Research under contract number DE-AC02-05CH11231.

## Data Availability

Genome assemblies from *A. baumannii* strains used for the initial bioinformatics analysis are described in **Supplemental Table S1**. A list of all TIGER and Islander-predicted prophages are in **Supplemental Table S2. Supplemental Table S3** contains annotations for prophage-specific proteins found in genome assemblies of NCBI phages that cluster with our TIGER/Islander-predicted phages. *Acinetobacter*-infecting phage accessions from NCBI are described in **Supplemental Table S4**. Primers used for this study are listed in **Supplemental Table S5**. Probes used to detect recombinant reads in Illumina sequencing data are listed in **Supplemental Table S6**. Individual phage capsid and tail measurements can be found in **Supplemental Table S7**.

## Notes

### Competing Interest Statement

The authors have declared no competing interest.

