## Supplemental Figures for "Discovery, induction, and screening of prophages in clinical *Acinetobacter baumannii* isolates"

### Supplemental Materials

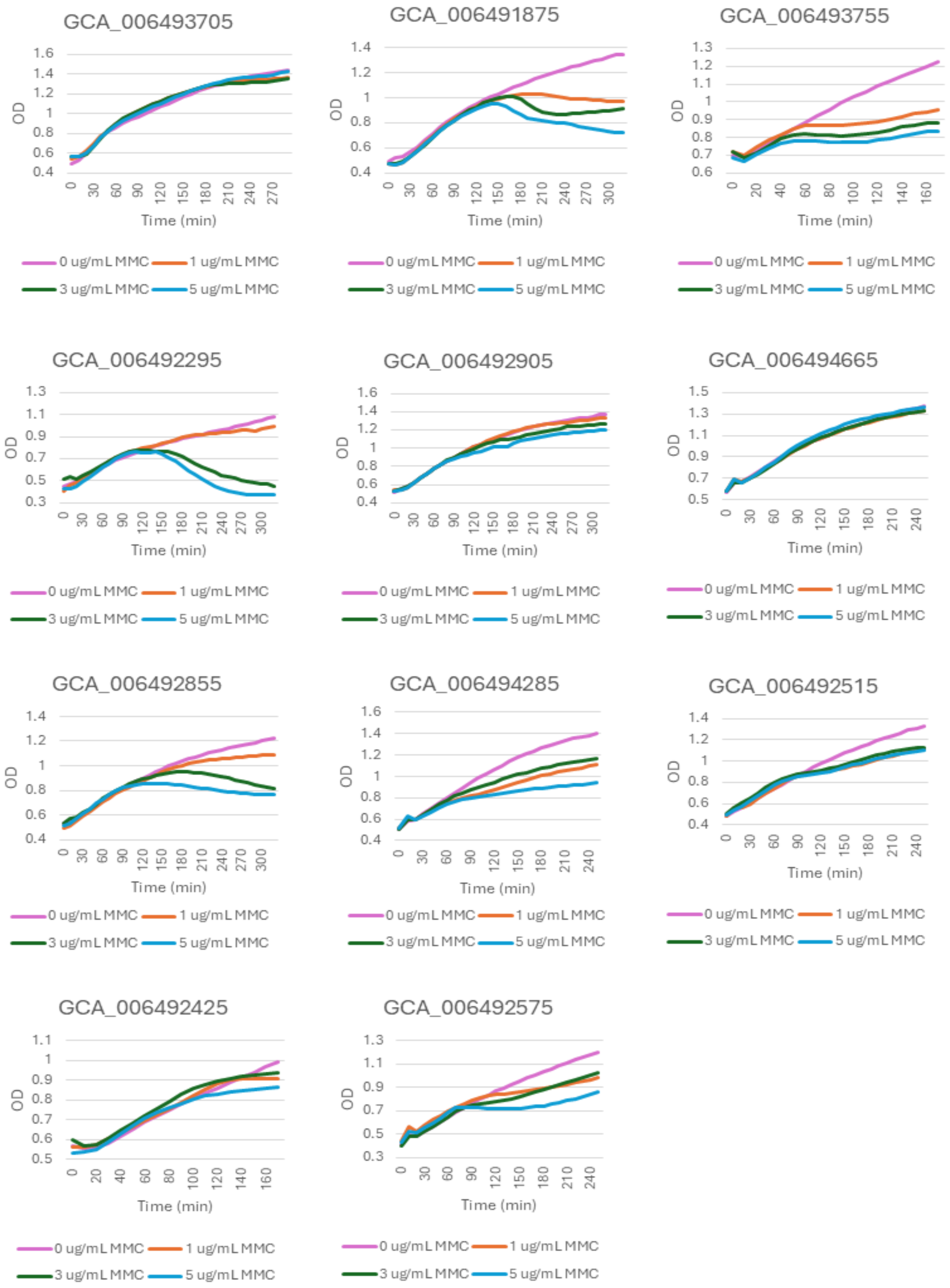


**Supplemental Figure S1. Growth curves of *Acinetobacter baumannii* strain induced with varying concentrations of mitomycin C (MMC).** Growth curves are the average of three technical replicates (wells) within the same 96-well plate. Strains where the final OD of the MMC-induced strains is markedly lower than the final OD of the uninduced strains were selected for further screening.

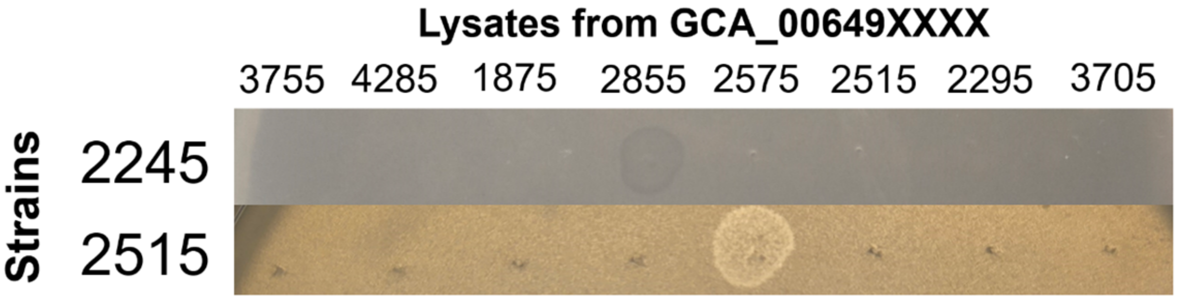


**Supplemental Figure S2. Lysates from two MMC-induced *Acinetobacter baumannii* strains can plaque on two different *A. baumannii* hosts.** 20 µL spots of lysate were placed on lawns of the respective *A. baumannii* strain on the left axis of the figure, and zones of clearing represent spots where bacterial growth was inhibited by the lysate.

**Supplemental Table S1. Table of *Acinetobacter baumannii* genome assemblies used in this study.** External IDs refer to those in the NCBI Nucleotide database.

**Supplemental Table S2. Table containing predicted prophages and their coordinates.** Only prophages with Phage1 categories are listed.

**Supplemental Table S3. Annotations of prophage genes in NCBI phages that cluster with TIGER/Islander-predicted prophage.**

**Supplemental Table S4. Table with all *Acinetobacter*-infecting phage genomes from NCBI used for analyses.**

**Supplemental Table S5. Primers for the amplification of attB and attP sites in *Acinetobacter baumannii* strains.**

**Supplemental Table S6. DNA sequences used as probes to quantify prophage excision and replication in sequencing data.** *attB* = unintegrated prophage, *attP* = excised and circularized phage, *attL* = start of integrated prophage, *attR* = end of integrated prophage.

**Supplemental Table S7. Capsid diameter measurements for phages induced from three *Acinetobacter baumannii* strains.** Measurements are given in nm. Note that GCA_006492575 has multiple types, with varying capsid and tail lengths.
